# Repurposed nystatin to inhibit SARS-CoV-2 and mutants in the GI tract

**DOI:** 10.1101/2021.10.19.464931

**Authors:** Eszter Virág, Dénes Seffer, Ágota Pénzes-Hűvös, Krisztina Varajti, Géza Hegedűs, István Jankovics, József Péter Pallos

## Abstract

The SARS-CoV-2 infections are considered as respiratory system diseases, mostly. In COVID-19, it might also be the infection of gastrointestinal (GI) tract too, especially at patients in severe clinical condition. SARS-CoV-2 can destroy the intestinal barrier, capable to spread into internal organs via blood and/or lymphatic circulation, and to cause serious damage there. Infected GI tract of COVID-19 patients is ideal environment for the coronavirus infection, replication and as virus reservoir might be the major source of pandemic reinfections, too. The process of virus budding is dependent on the host cell lipid rafts containing membrane-sterols, mainly cholesterol. The viral envelope may be challenged by polyene antibiotics, such as nystatin, which has strong affinity to sterols. Nystatin may block the establishment of the virus-host cell connection, too. In this study, the nystatin was investigated, as antiviral agent to SARS-CoV-2. We demonstrated by tests in Vero E6 cell based cytopathic assay, nystatin blocked the replication of SARS-CoV-2 in concentration 62.5 μg/ml (IC50) at Wuhan and British mutant strains. No efficient SARS-CoV-2 antiviral agent is known so far to alleviate pandemic, to disinfect GI tract, where vaccines might have limited effect, only. Nystatin might be the first one with emergency use authorization, either, as a safe and efficient non-systemic antiviral drug, with well-established use, since decades.

## Introduction

The severe acute respiratory syndrome coronavirus 2 (SARS-CoV-2) is originated from Wuhan in late 2019 and resulted in pandemic with crisis in public health and in World economy. More than 220 million cases and 4.5 million deaths have been registered by the Center for Systems Science and Engineering at John Hopkins University, in the beginning of September 2021. This virus belongs to the *Betacoronavirus* genus of *Coronaviridae* family of enveloped, single-positive-stranded RNA viruses, which may cause infection in lung and the gastrointestinal tract (GIT), as well. They are sensitive to bile-acids, to intestinal proteases, which may alter membrane lipid distribution and function of proteins attached to lipid rafts, making them more ineffective (1–3). SARS-CoV-2 causes mainly upper respiratory tract symptoms however the virus presence in the GIT is remarkable. GIT symptoms are also well known in 2% to 79% (4–8) in COVID-19 and the GIT symptoms were associated with the severity of the disease (9). A meta-analysis of 21 studies with 5285 patients draws attention to COVID-19 to be more serious in patients with GIT symptoms exploring the relationship between GIT symptoms and the severity of COVID-19 (10). Another meta-analysis of publications found that stool samples from 48.1% of patients tested positive for virus RNA and stool samples from 70.3% of these patients tested positive for virus RNA even after respiratory specimens tested negative (11). Diarrhea has been listed as a symptom of COVID-19 in the guideline of the Centers for Disease Control and Prevention since 22 February 2021 (12).

First the American College of Gastroenterology has drawn attention to “patients with new-onset digestive symptoms after a possible COVID-19 contact should be suspected for the illness, even in the absence of cough, shortness of breath, sore throat…” (13). This important guideline, now is well strengthened by the tool described here, to cure this illness, in GIT importantly. In spring of 2021 Leal et al. reported on GIT symptoms present in 30% of the European patients, among which diarrhea was the most frequent with almost 18% incidence (14). Similar study on the American population resulted in 22.4% incidence. (9). Gastrointestinal mucosal damage (degeneration or necrosis) (15), bleeding (7, 16, 17), edema (18–20) were present in severe COVID-19 cases suggesting a direct effect of the virus on GIT cells. Elevated liver enzymes was observed during hospitalization and progression to severe liver injury have also been noted in COVID-19 patients. The severe decompensated liver disease increased the severity of COVID-19 and vice versa (21). Despite the fact of above-mentioned clinical symptoms only a few study reports on the investigation of GIT tissue samples (22). Papoutsis et al. identified and characterized mutational variations of SARS-CoV-2 by enrichment next-generation sequencing (NGS) from stool samples suggesting that the gut is an ideal environment for the virus, where it may also mutate (23). The role of gut during the SARS-CoV-2 pathogenesis was proved by detecting viable and infectious viruses from stool samples (24–26). Several studies report on the isolation of virulent virus after the resolution of GIT symptoms even when it was not detectable from respiratory samples. This phenomenon based the ability of virus replication in GIT which may contribute to long-term consequences after the disease (27, 28). Multisystem inflammatory syndrome in children is thought to driven by zonulin-dependent loss of gut mucosal barrier which is the consequence of the prolonged presence of SARS-CoV-2 in the GIT (29).

All these data highlight the importance and the urgent need of the local antiviral treatment of the GIT for which we are looking for a solution.

The mechanism of the SARS-CoV-2 viral infection is investigated thoroughly. The key factor for viral entry and life cycle is the cholesterol in the human plasma membrane (30). Enveloped viruses primarily engage plasma membrane fusion or endocytosis for entering the host cell (31). Lipid raft domains are involved in this process and serve as a platform for docking the viruses to the host cell. Li et al. found that lipid rafts may contribute to SARS-CoV-2 infection during the replication process in Vero E6 cells (32). The augmented cholesterol/ fatty acid ratio of lipid rafts enhances the fusion of coronaviruses to the host cells while the decreased ratio blocks this process (33). Meher et al. reported on the SARS-CoV-2 infection rate and binding affinity was increased by raising the cholesterol level in human plasma membranes (30). Wang et al. proved that decreasing membrane cholesterol inhibits SARS-CoV-2 entry (34). Sanders et al. found that in the case of SARS-CoV-2, cholesterol is essential for the spike-mediated fusion, and for the pathological multinucleated cell (syncytia) formation as well (35). Wang et al. showed that loading cells with cholesterol from serum, using a cholesterol transport protein (apoE) enhanced the entry and the infectivity of the SARS-CoV-2 spike protein pseudotyped retro virus (36). Cholesterol rich SARS-CoV-2 entry sites showed almost twice the total endocytic entry sites. Additionally, they found that in virus-producing cells, the cholesterol optimally positions a protease enzyme, furin for priming the virus. Inhibiting cholesterol transport has been found to inhibit SARS-CoV-2 replication in late endosomes/lysosomes (37). Also, depletion of the available cholesterol of plasma membranes inhibits the virus-membrane fusion (34).

In the mechanism of receptor mediated cell entry, the role of angiotensin converting enzyme II (ACE2) was proved (38, 39). Organs such as of the respiratory tract, GIT, bile duct and liver have a high expression of ACE2 receptors making them a specified target for SARS-CoV-2 (21). A single-cell RNA-Seq analysis showed that the ACE2-positive-cell ratio in digestive tract organs was significantly higher compared to the lung in COVID-19 patients (40) and the bile duct endothelial cells expressed higher quantities of ACE2 receptor than the liver endothelial cells (41). This data may correlate with that ACE2 is an important regulator of intestinal inflammation (42). Toełzer et al. report on the cryo–electron microscopy structure of SARS-CoV-2 spike (S) glycoprotein (S-protein) revealing that the receptor-binding domains tightly bind the essential free fatty acid linoleic acid (LA) in three composite binding pockets. LA binding stabilizes a locked S conformation, resulting in reduced ACE2 interaction in vitro (43).

The possible role of high-density lipoprotein (HDL) particles during SARS-CoV-2 infection was also studied. Hu et al. hypothesized that because of the immunomodulatory effects of HDL-cholesterol (44, 45), it may be involved in the immune cell regulation, thus resulting in the decreased HDL levels in COVID-19 patients (46). SR-B1 is a cell-surface HDL receptor that mediates the selective uptake of receptor-bound HDL particles therefore SR-B1 has a critical role in hepatitis C virus entry (47). Based on this mechanism a potential role of SR-B1 in SARS-CoV-2 infection was raised. Wei et al. found that the SR-B1 overexpression in Vero E6 cells increased SARS-CoV-2 infection. Additionally, the S-protein is bound to cholesterol with high S1 subunit affinity. However, S-protein did not bind to ApoA1 protein (48), main component of HDL which enhanced the entry of SARS-CoV-2. These data suggested that HDL might form a bridge between SARS-CoV-2 and SR-B1 during viral entry (48).

The role of HDL in COVID-19 was manifested in the change of HDL plasma concentration and proteome composition and functionality as severe patients were associated with HDL dysfunction manifesting in aggravated inflammatory endothelial conditions (49, 50). Another evidence for lipid involvement in COVID-19 pathogenesis was suggested by Nardacci et al. They observed lipid droplets accumulation in cells during SARS-CoV-2 infection, both in vitro and in lungs of COVID-19 patients (51). Correlations between SARS-CoV-2, cholesterol, HDL, lipid rafts and fatty acids are summarized in Figure. 1.

**Figure 1.**
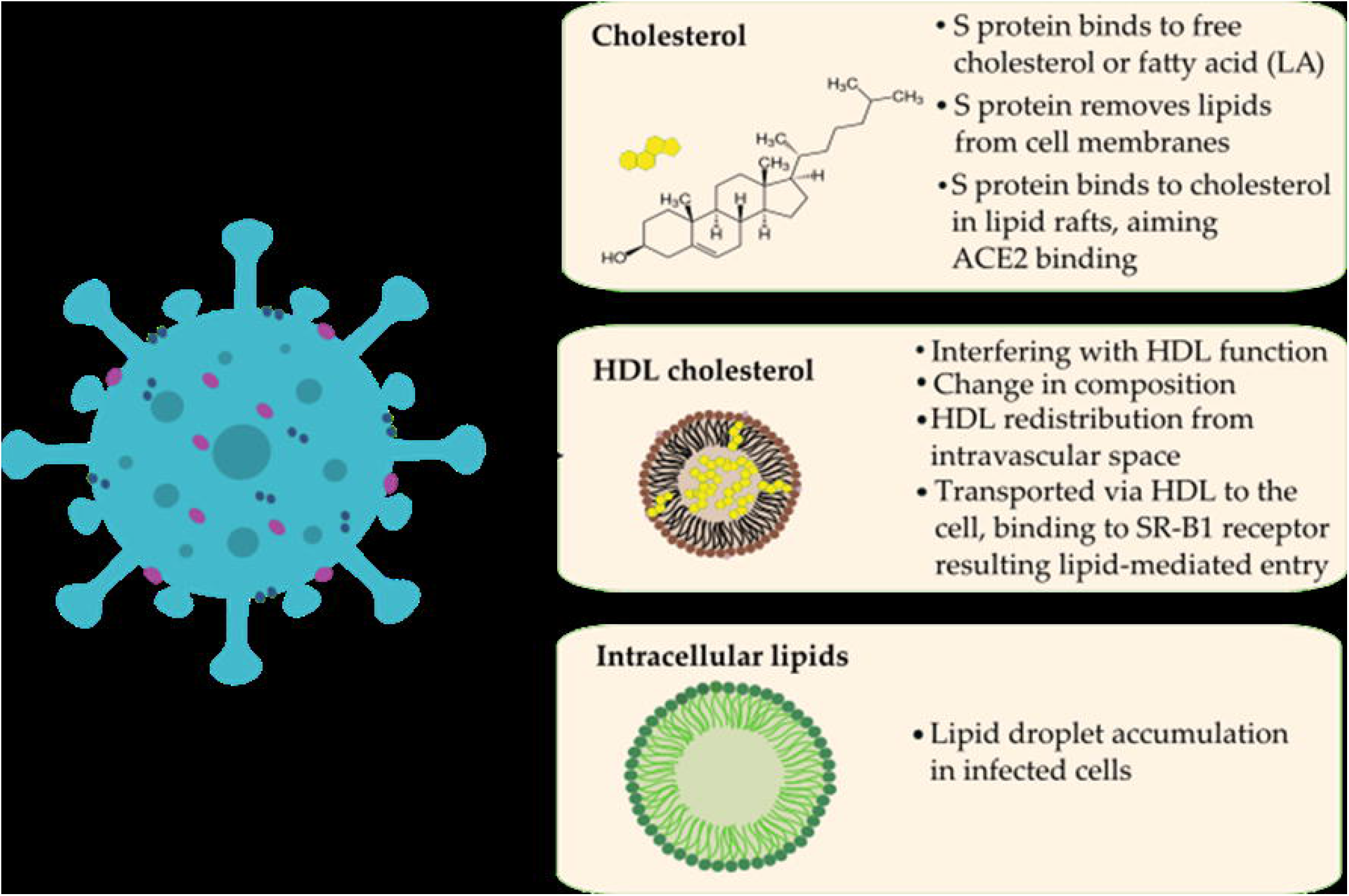
Possible correlations between SARS-CoV-2 (left) and lipids (LA) / HDL-cholesterol (right) during pathomechanism based on the literature.

Compounds affecting cholesterol were shown to interfere with viral infectivity at different stages of viral replication. For example, cholesterol depletion proved to disrupt the HIV-1 virion membrane (52). Baglivo et al. report on natural-based substances (cyclodextrins and phytosterols) that were used to reach membrane destabilization through cholesterol reduction and disturbed the process of SARS-CoV-2 entry into the host cells (53). Another cholesterol-lowering agent, fenofibrate recently has been proved to be effective in the reduction of SARS-CoV-2 infection *in vitro*. Treatments for reducing the plasma cholesterol levels (i.e., by the use of statins) are also suggested by healthcare professionals for a long while in COVID-19 care (54).

Nystatin is a polyene macrolide antibiotic whose mechanism of action is to bind to sterols and forming ion channels in the fungal membrane leading in this way to leakage and cell death. This compound possesses a high affinity to bind to the ergosterol of the fungal plasma membrane, but the binding to human cholesterol is also significant (55, 56). Possible interactions between nystatin and cholesterol are presented in Figure. 2. Nystatin is not appreciably absorbed from the mucous membranes of GIT, thus has no systemic effect from oral formulations and therefore can be used to treat fungal infections of the GIT. Consequently, it may modify the cell surface through lipid rafts ‘floating’ on the cell surface, binding to their cholesterol content to prevent the SARS-CoV-2 viral particle from entering the cell (57, 58). This may require a high concentration of active ingredient that could be systemically problematic based on its toxicity, but not in local application. It is also reported in the literature that polyene macrolides, such as nystatin, may also be able to alter the fat metabolism of mammalian cells, thereby interfering with the replication of the virus (32). In any case, nystatin-based pharmaceuticals are already indicated for use as complementary therapy as preventing candidiasis in patients assigned to antibiotic therapy.

**Figure 2.**
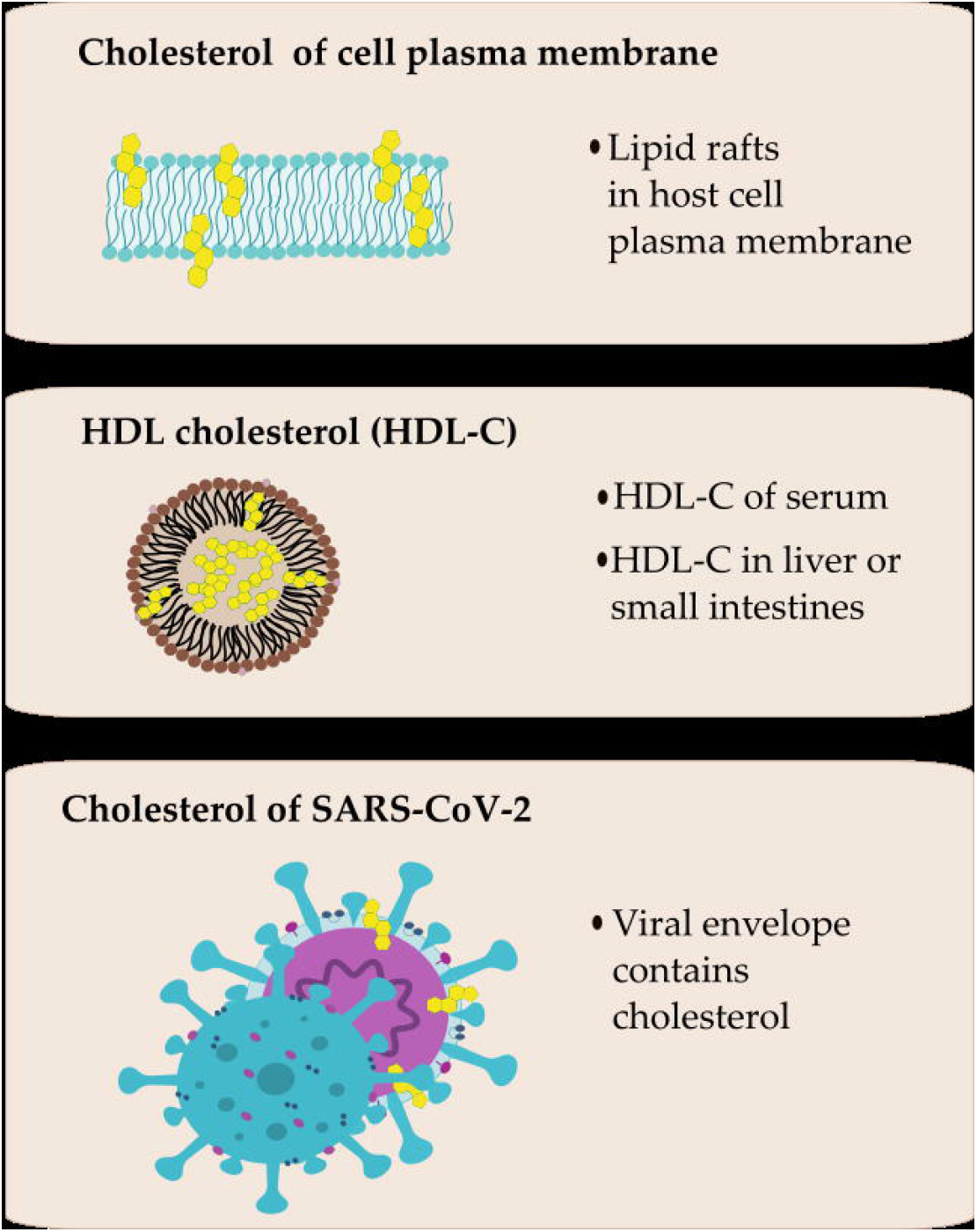
Possible interactions between nystatin (left) and cholesterol of different origin (right).

Nystatin had been previously examined regarding to its potential antiviral effect in HIV-1 infection model in H9 cells by Selvam et al (59). Nystatin A inhibited viral replication effectively, in certain concentrations that did not affect cell viability. The results suggest that Nystatin A merited attention as an antiviral drug for the treatment of HIV-1 infection. Nystatin in the antiviral indication alone was used against HIV in a clinical trial also (60). Nystatin had also been proved to be effective in reducing the cytopathic effect manifesting in cell-cell fusion during infection of SARS-CoV-2 pseudovirus (35). In addition to the antifungal and antiviral effects of nystatin, its immunomodulatory effect was also proved (61, 62).

We recommend that nystatin may be a safe and effective agent in the treatment of infections caused by enveloped viruses, including COVID-19. Since this drug affects the virion structure as well and is not absorbable in the GIT, the development of nystatin resistance has less chance and nystatin may has a direct antiviral effect on viruses during infection, while might have only negligible effect on host tissues. To verify the antiviral activity of nystatin against SARS-CoV-2 *in vitro*, the virus inhibitory effect of nystatin was investigated performing SARS-CoV-2 antiviral assay on Vero E6 cells according to the method of Manenti et al. and Amanat et al. (63, 64).

## Materials and Methods

### Materials

Nystatin was used in pharmaceutical grade. 50.000 μg/ml stock solution in DMSO diluted 100-fold in Dulbecco’s Modified Eagle’s Medium (DMEM)-high glucose supplemented with 2 mM L-Glutamine, 100 units/ml penicillin-streptomycin mixture without FBS.

### Cell cultures

Vero E6 cells were purchased from Veterinary Diagnostic Directorate of the National Food Chain Safety Office, Hungary. The cells were cultured in Dulbecco’s Modified Eagle’s Medium (DMEM)-high glucose (Sigma) supplemented with 2mM L-Glutamine (Sigma), 100 units/ml penicillin-streptomycin mixture (Sigma), and 10% of FBS, at 37°C, in a 5% CO_2_ humidified incubator. Adherent sub-confluent cell monolayers were prepared in growth medium DMEM high glucose containing 2% FBS in T175 flasks or 96-well plates for propagation or titration and neutralization tests of SARS-CoV-2.

### Viral growth in cell culture

Vero E6 cells were seeded in T175 flasks at a density of 1 × 10^6^ cells/ml. After 18 to 20 hours, the sub-confluent cell monolayer was washed twice with sterile Dulbeccos’s phosphate buffered saline (DPBS). After removal of the DPBS, the cells were infected with 3.5 ml of DMEM 2% FBS containing the virus at a multiplicity of infection of 0.01. After 1 hour of incubation at 37°C in a humidified atmosphere with 5% CO_2_, 50 ml of DMEM containing 2% FBS were added to the cells. The flasks were daily observed and the virus was harvested when 80%-90% of the cells manifest cytopathic effect (CPE).

### Virus and titration

SARS CoV-2 virus strains (Wuhan and British mutants, CMC-1 and VEVE) were obtained from the Complex Medical Centre, Budapest, Hungary. For detailed informations of strains, see Supplementary material 1. The virus was titrated in serial dilutions of 1log to 11log (10^−1^ to 10^−11^) to obtain 50% tissue culture infective dose (TCID50) on 96-well culture plates of Vero E6 cells. The plates were observed daily for a total of 4 days for the presence of CPE by means of an inverted optical microscope. The end-point titer were calculated according to the Reed & Muench method based on eight replicates for titration (65).

### Treatment of virus

Two-fold serial dilutions of nystatin, starting from 500 μg/ml were prepared. All dilutions contained 1% DMSO. The dilutions were mixed with an equal volume of viral solution containing 100 TCID50 of SARS-CoV-2 in Dulbecco’s Modified Eagle’s Medium (DMEM)-high glucose supplemented with 2 mM L-Glutamine, 100 units/ml penicillin-streptomycin mixture. The nystatin-virus mixture was incubated for 30 min. at 37°C in a humidified atmosphere with 5% CO_2_.

### Layout of SARS-CoV-2 antiviral assay

96 well plates (Figure 3) were prepared according to the following. A1-H12 wells contained Vero E6 cells (0.2 × 10^6^ cell number per well). Column 12 contained the control cells (CC): cells without virus and nystatin treatment. Column 11 contained the virus control (CV): 100 TCID50 of SARS-CoV-2 without nystatin treatment. Column A1-D10 contained virus treated with nystatin dilution series. 100 μl of the mixture at each dilution of nystatin was added in quadruplicate (A1-D1, A2-D2, etc.). Column E1-H10 contained cells treated with the nystatin dilution series in quadruplicate without virus (E1-H1, E2-H2, etc.). The bisecting dilution series of nystatin concentration was 500 μg/ml - 0.97 μg/ml. The plates were incubated for 4 days at 37°C in a humidified atmosphere with 5% CO_2_.

**Figure 3.**
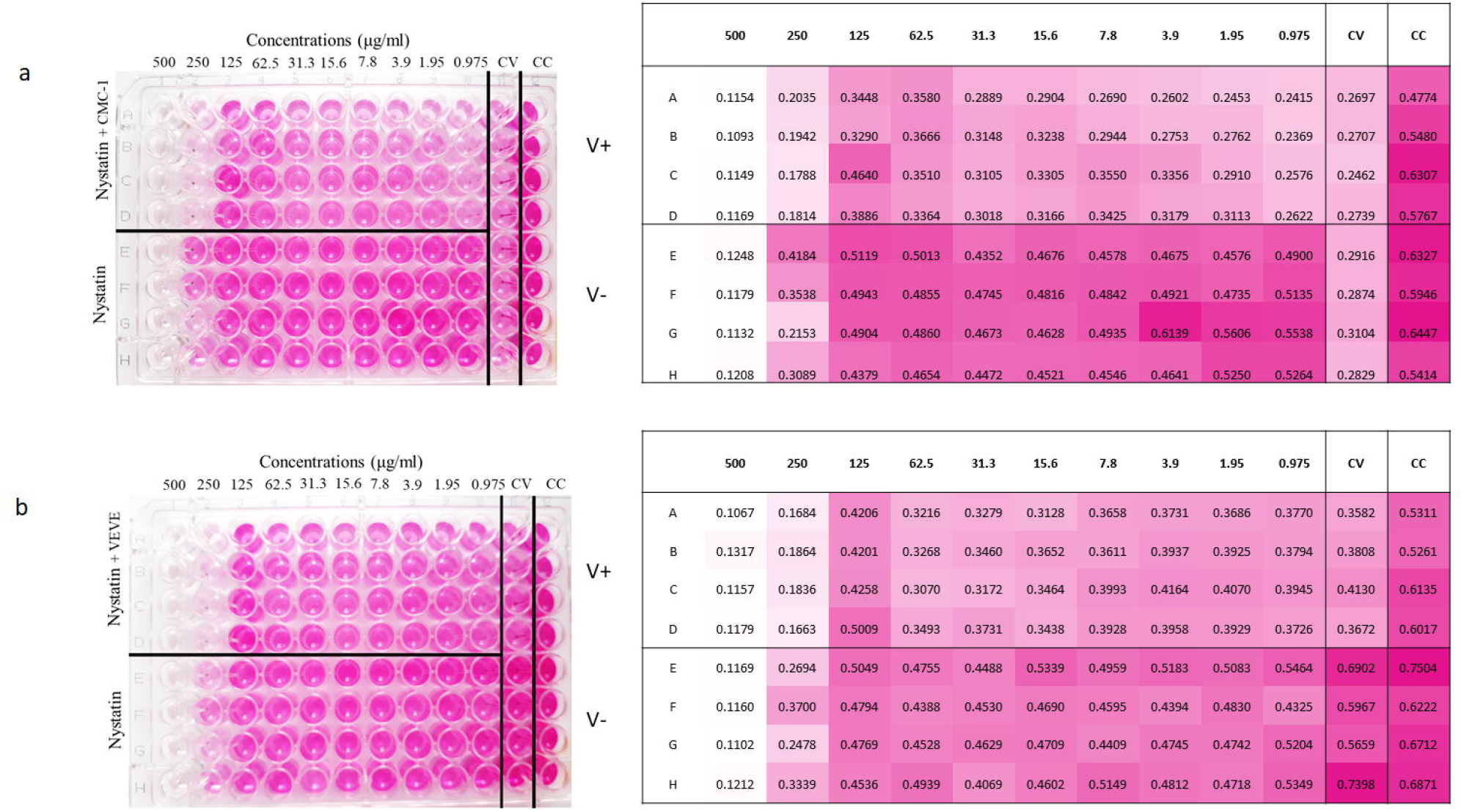
Colorimetric read-out of SARS-CoV-2 antiviral assay of nystatin. The cell plate contained a semi-confluent Vero E6 monolayer with SARS-CoV-2 infections of Wuhan strain, CMC-1 (a) and British strain, VEVE (b). Abbreviations: virus control (CV), cell control (CC), cells infected with SARS-COV-2 (V+), cells without virus infection (V-).

### Colorimetric read-out of SARS-CoV-2 antiviral assay

After 3 days of incubation, the supernatant of each plate was carefully discarded and 100 μl of a sterile DPBS solution containing 0.02% neutral red was added to each well. After 1 hour of incubation at room temperature, the neutral red (Renal) solution was discarded and the cell monolayer was washed twice with sterile DPBS containing 0.05% Tween 20. After the second incubation, the DPBS was carefully removed from each well; then, 100 μl of a lysis solution (made up of 50 parts of absolute ethanol, 49 parts of MilliQ and 1 part of glacial acetic acid) was added to each well. Plates were incubated for 15 minutes at room temperature and then read by the spectrophotometer infinite F50 at 492 nm and by using Tecans iControl Software. The plates were inspected by an inverted optical microscope before colorimetric read-out.

## Results

### Antiviral effects of nystatin on the replication of SARS-CoV-2 in cell culture

To confirm that nystatin inhibit SARS-CoV-2 replication, Vero E6 cell culture were infected with two mutants of SARS-CoV-2 pretreated with nystatin *in vitro*. SARS-CoV-2-early Wuhan and British mutants were treated with increasing concentrations of nystatin, and protection from cytopathic effects (CPE) observed visually and with colorimetric read-out. Nystatin showed a reduction of the viral CPE dose dependently (Figure 3). After the cells were stained with neutral red, their optical density at 492 nm was measured (Figure 3a). At 96 h post infection, cells were examined with an inverted optical microscope (magnification, x100). The cell control (CC) refers to cells without compound treatment and virus infection (Figure 4a). Vero E6 cells after 96 h of SARS-CoV-2 infection are shown on the Figure 4c as the virus control (CV). 62.5 μg/ml nystatin treated SARS-CoV-2-infected cells are shown in Figure 4b. The results showed that nystatin inhibited SARS-CoV-2 infection, with an effective concentration (EC) of 62.5 μg/ml against Wuhan CMC-1 and British mutant VEVE strains (Figure 5). To evaluate the cytotoxicity of nystatin to cells, Vero E6 cells were treated with different concentrations of nystatin up to 500 μg/ml. Taking into account the toxic effect, the concentration at which nystatin inhibited the virus growth, the EC was determined. We found that nystatin was only slightly toxic up to 250 μg/ml

**Figure 4.**
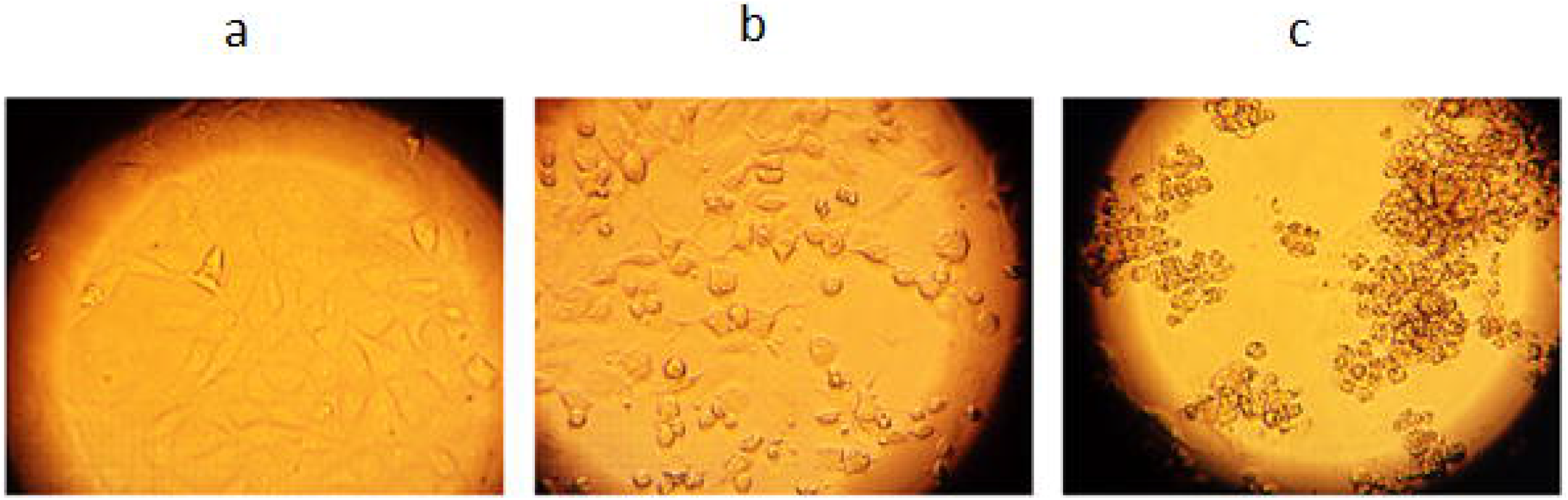
Validation of viable cells by inverted optical microscope. Cell control, CC (a); cells with SARS-CoV-2 Wuhan strain treated by 62.5 μg/ml nystatin (b); virus control, CV (c).

**Figure 5.**
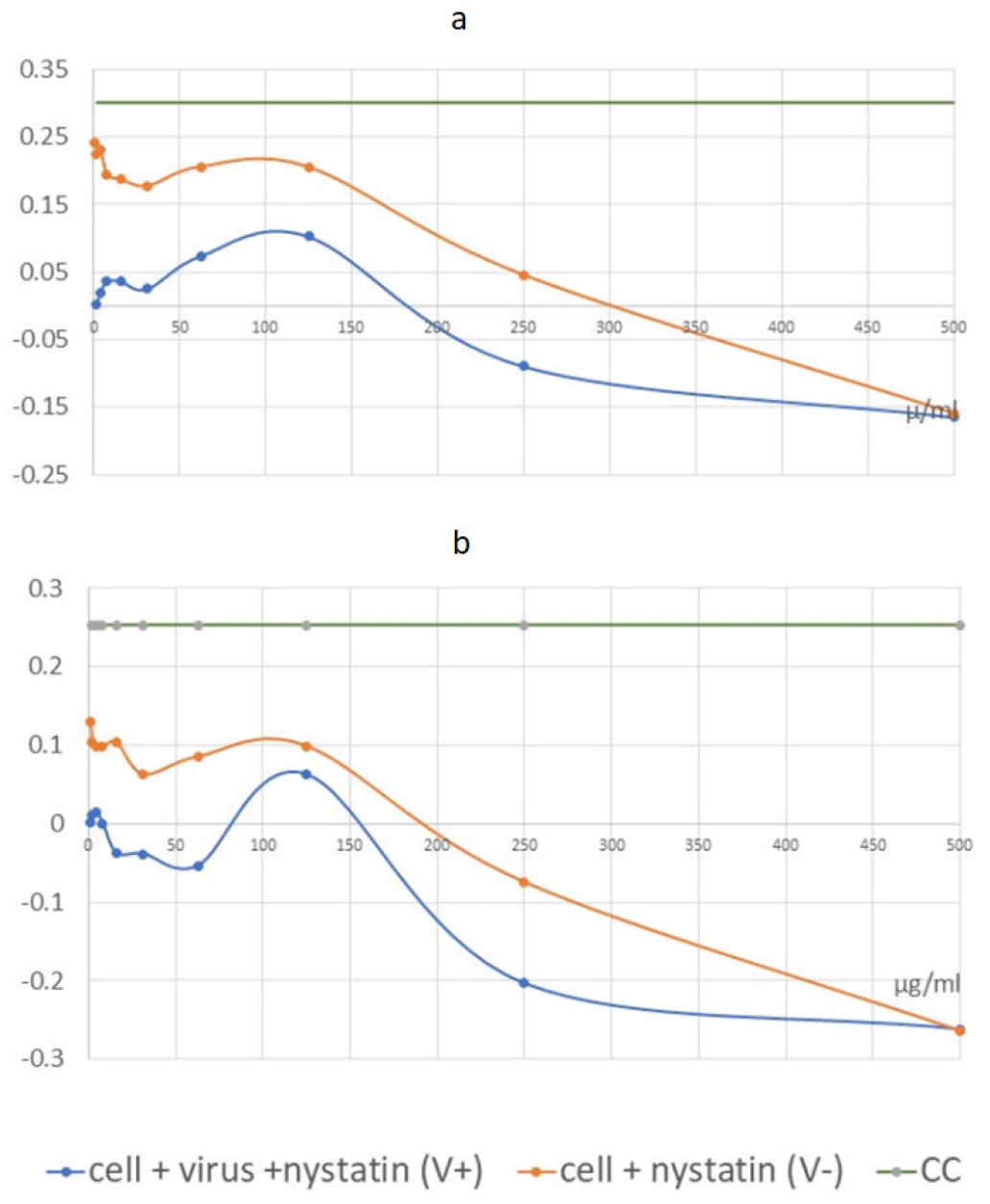
Graphical illustration of the corrected absorbance values. IC50 of nystatin may be detected at 62.5 μg/ml for the Wuhan strain, CMC-1 (a) and 125 μg/ml for the British mutant, VEVE (b).

### Calculation of dose response to nystatin

Values of the measured absorbance are proportional to the amount of living cells. To evaluate the plates, the absorbance values of the four replicates were averaged. CV was then extracted from all values to obtain corrected absorbance (Figure 5). In the case of CV, almost complete cell death was observed. In the case of CC, the cell layer confluently overgrown the wells. Where the corrected absorbance was positive greater than cutoff (63) – OD (CC) /2 – there were cells that appeared alive confirmed by microscope (Figure 4b).

Based on our investigation, it can be determined that treatment with nystatin up to 125 μg/ml was not cytotoxic, but at all concentrations reduced the live cell concentration compared to the cell control (CC) to an extent that was considered constant independent of the concentration used. Nystatin treatment has been shown to be effective against CMC-1 (Figure 5a) and VEVE (Figure 5b) SARS-CoV-2 strains at concentrations of 62.5 and 125 μg / ml, respectively. Since the accurate antiviral IC50 could not be determined from reason of nystatin-host cell effect, the EC value of nystatin was modelled mathematically (Figure 6).

**Figure 6.**
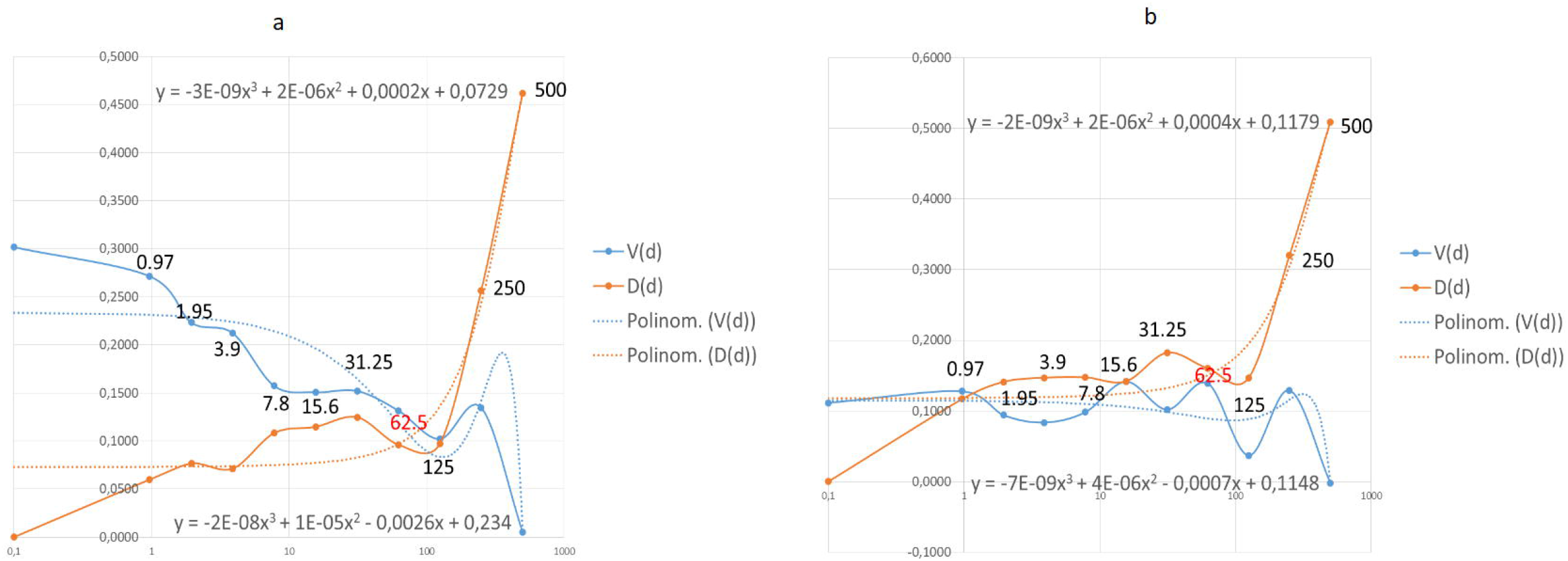
Effective dose (EC) of nystatin was calculated by data analysis with mathematical functions. Points where the functions are stable are the 62.5 in both Wuhan (a) and British (b) mutant strains. The functions are not stable at the grater points therefore the effective and safe dose is 62.5 μg/ml in both case. The values shown in the diagram are the nystatin doses correlated to function points.

### Mathematical data analysis of EC value of nystatin

EC value of nystatin was modelled by a mathematical data analysis (Figure 6). Functions were generated such as: (i) V(d) a dose-dependent viral function, where (d) is an independent variable. This function determines how much the agent changes the number of cells by the effect of the dose on the number of viruses, which reduces the number of cells by a viral function definition. Thus, it describes the change in cell number by the virus in the dose function. (ii) D(d) is a function that determines which cell number definition is affected by the dose and directly by the agent on the cell number. To get the D(d) function need to know A, that is the average of CC: 0.5808.

Describing by a function, there is present both virus and agent: A-V(d) -D(d) this means A-D on the plate rows. To determine D the following definition can be determined:

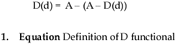

A polynomial of degree 3 functional can be fitted to both. It can be seen that at high doses the destructive effect of the virus decreases, while with increasing dose the D function reduces the number of cells increasingly. The optimum must be found where the dose level of the agent is still safe. The closest dose value to the intersection (Figure 6a) of the trend line is 62.5. 125 can also be considered as a close point, mathematically, and may be a good solution, but it is the limit of a drastic change in value for both trend lines, so it is less safe to use this dose (Figure 6a). The same phenomenon can be seen in Figure 6b. The last point where the functionals shown stability is 62.5. This is the concentration of which nystatin shown the highest efficacy with low toxicity.

The IC50 calculated according to the standard protocol (Figure 5) was not exact to estimate the antiviral dosage of nystatin in this test system. The values of effective antiviral dose therefore was based on mathematical modelling suggesting 62.5 μg/ml in both case.

In general use, nystatin dose is 100 mg-600 mg daily. Since nystatin is a non-absorbable drug this means a dilution in 5-6 liter GIT juice daily (66), approximately. In this experiment we determined the EC under the maximum daily dose of nystatin.

## Discussion

During the SARS-CoV-2 pandemic period of time, - as the virus’ name Severe Acute Respiratory Syndrome shows accordingly-COVID-19 is considered mainly as a new disease in pulmonology, and viral infection, invasion of GIT are oversight and undervalued, equally. However, the increasing concentration of excreted viral fragments in wastewater, as shown by tests data, might predict the soon increase in community infection rate, days, or weeks earlier than the number of clinical cases with symptoms of COVID-19 are set on, in communities (67). The web search analytics as infodemiology study for abdominal symptoms in COVID-19 reflects the early sign of infection (diarrhea) presenting the involvement, i.e. the viral infection of gastrointestinal tract of patients, at the same time (68, 69). In addition to the additional GIT manifested clinical cases, notable number of asymptomatic and presymptomatic virus carriers are also to be considered as uncontrolled role-players in pandemic (70). The pandemic of SARS-CoV-2 requires an intensive search for new antiviral agents. Many compounds are currently being investigated for their efficacy in COVID-19 therapy. Considering the severity of COVID-19 there is a need to find effective but less toxic drugs for a rapid treatment. Therefore the possibility of non-systemic therapy came to the fore. In this report the antiviral effect of nystatin antibiotic was investigated against Wuhan and British mutant strains of SARS-CoV-2 tested on Vero E6 cells. The presented results demonstrate that nystatin at a non-cytotoxic dose may protect cells against infection with SARS-CoV-2. This protection is associated with inhibition of virus replication that was manifested by an augmented living cell rate (Figure 3).

Nystatin is a polyene macrolide antibiotic of which the broad-spectrum antifungal activity and low fungal resistance are characteristics (71–75). Nystatin is not or slightly absorbed (76, 77) and non-toxic (77) allowing its application in the GIT. It is used worldwide since 1961, in different pharmaceutical formulations and dosage forms in antifungal indications, including prophylaxis. It might be a safe agent as the annual number of reports on adverse reaction counts less than 100 per year in average (78). The antiviral effect of polyene macrolide antibiotics against a variety of lipid-enveloped RNA and DNA viruses was proved (59, 79, 80). Nystatin bind irreversibly with sterols (81, 82). This is a direct structural and non-metabolic effect that consequence is the low resistance rate (83, 84). The binding of nystatin to cholesterol in cell membranes causes changes in plasma membrane fluidity and cell permeability therefore decreases infectivity of lipid-enveloped viruses. The drug-sterol interaction is a well-known phenomenon that has already been demonstrated in our earlier studies on the macrolide antibiotic primycin (84–90) where the membrane disorganization was due to the ergosterol-primycin bound stabilized by hydrogen bounds (87).

The inactivation of SARS-CoV-2 is dependent on the dose of nystatin: the accurate dose of inhibition is probably due to a complex effect that is strongly depend on the binding action of host-cell membrane and/or viral envelope cholesterol ratio. Similar effect of nystatin was described against HIV-1 (59). Interestingly, if viral envelope cholesterol was reduced the infectivity of SARS-CoV-2 decreased while depletion of cell membrane cholesterol had no effect on the virus infectivity (35). Thus, we suppose that the envelope cholesterol/nystatin molecule ratio may influence that drug efficacy. In dose determination we had to take into account a slight restrictive effect of nystatin (Figure 3) on test-cells connected to the inhibition effect of virus. Based on this 62.5 μg/ml nystatin concentration showed to be the effective dose which improved the infected Vero E6 cells’ viability.

Considering potential therapeutic usage, the properties of non-toxic and LADME - which represents the main pharmacological characteristics of any pharmaceutical product - of nystatin should be interpreted. Liberation, Absorption, Distribution and Metabolism steps are hardly explainable due to the poor water solubility and lack of absorption of nystatin. Liberalization and distribution are limited to the GIT, metabolism is restricted to the interaction with the gastrointestinal juice and its fecal elimination is known. Based on the well-established antimicrobial indication, the daily dose of nystatin is 100-600 mg which is continuously diluted or concentrated depending on secretion or reabsorption of the GIT segments. In this way, the total dilution may be around 5-6 liter in GIT juice daily **(66)**, thus the determined effective antiviral concentration is only about the half of the maximum allowed daily concentration of nystatin. This means there is no reason to change the well-established dosage regimen in a potential SARS-CoV-2 GIT disinfection therapy either.

In our opinion, GIT disinfection pharmaceutical products may contribute to a step forward in overcoming the pandemic on different levels. Individuals (patients with GI symptoms) shall benefit sooner recovery and less severe or fatal, recurrent, and postcovid cases may be experienced. For communities, one may expect a smaller number of asymptomatic carriers, making lockdown periods more effective and contribute to a better control on pandemic. From regulatory point of view, the issue of emergency use market authorization to efficient, properly documented (i.e. cytopathic assay evidenced SARS-CoV-2 antiviral) nystatin formulations shall be considered.

To summarize. Considering the urgency of the ongoing COVID-19 pandemic, detection of various new mutant strains and the future potential re-emergence of novel coronaviruses, repurposing of authorized drugs such as nystatin could be worthy of attention. Vaccines mainly have systemic effect, they do not neutralize viruses in GIT, one of the general sources of viral infection. Our results suggest that a non-systemic nystatin therapy – based on virus inactivation-, in cytopathic assay proven formulated drugs may provide a solution for COVID-19 therapy through GIT disinfection. The use of this safe drug in preventive, prophylactic and therapeutic treatment might offer healthcare and economic advantages and benefits additional to the worldwide vaccination.

## Acknowledgments

Dr. Márton Milánkovits, Hungarian National Institute of Genitourinary Medicine Foundation, Hungary, drew our attention to the possible antiviral healing effect of nystatin. Based on this, we conducted our research, which we can now present to the scientific audience. We express our thanks to Barbara Kutasy, Hungarian University of Agriculture and Life Sciences, Hungary, for graphical illustration.

